# Wind and route choice affect performance of bees flying above versus within a cluttered obstacle field

**DOI:** 10.1101/2021.10.08.463704

**Authors:** Nicholas P. Burnett, Marc A. Badger, Stacey A. Combes

## Abstract

Bees flying through natural landscapes encounter physical challenges, such as wind and cluttered vegetation. The influence of these factors on the flight performance of bees remains unknown. We analyzed 548 videos of wild-caught honeybees (*Apis mellifera*) flying through an enclosure containing a field of vertical obstacles that bees could fly within (through open corridors, without maneuvering) or above. We examined how obstacle field height, wind presence and direction (headwinds or tailwinds) affected altitude, ground speed, and side-to-side casting (lateral excursions) of bees. When obstacle fields were short, bees flew at altitudes near the midpoint between the tunnel floor and ceiling. When obstacle fields approached or exceeded this midpoint, bees typically, but not always, increased their altitudes to fly over the obstacles. Bees that flew above the obstacle fields exhibited 40% faster ground speeds and 36% larger lateral excursions than bees that flew within the obstacle fields, likely due to the visual feedback from obstacles and narrow space available within the obstacle field. Wind had a strong effect on ground speed and lateral excursions, but not altitude. Bees flew 12-19% faster in tailwinds than in the other wind conditions, but their lateral excursions were 19% larger in any wind, regardless of its direction, than in still air. Our results show that bees flying through complex environments display flexible flight behaviors (e.g., flying above versus within obstacles), which affect flight performance. Similar choices in natural landscapes could have broad implications for foraging efficiency, pollination, and mortality in wild bees.

## INTRODUCTION

Nectivorous insects such as bees (Hymenoptera: Apoidea) provide important ecosystem services by pollinating wild plants and crops, and these services are intimately linked to their ability to successfully move through complex habitats while foraging for floral resources [1]. Bees typically move between their nest site and food sources by flying, sometimes over distances of several kilometers [2], yet many physical features of bees’ habitats can make flight a demanding, dangerous, and energetically expensive task [3–5]. Bees regularly fly through environments containing unpredictable winds and cluttered vegetation, each of which can pose distinct challenges to flight [5–8]. Furthermore, the heterogeneity of natural landscapes results in multiple route options for bees, so the unique behavioral choices made by individual bees can dictate the microhabitat (e.g., wind, clutter) that bees encounter, and thus the specific flight challenges that they must overcome [9]. Although bees foraging in wind and around vegetation is an everyday sight, we know surprisingly little about the effects of these habitat features on the flight behavior and performance of bees.

Cluttered vegetation can pose mechanical challenges to flying bees, but at the same time it provides visual landmarks that help bees navigate their environment [9–12]. The structure of vegetation can vary in many ways, including plant density, plant height, leaf size, and branch size [13–15]. Bees find navigable paths through clutter by using both brightness gradients within gaps (where brightness increases with gap size) [11] and optic flow, which is the apparent motion of the landscape moving past a bee’s eyes [16–19]. In particular, optic flow helps bees navigate clutter under a variety of conditions because it depends on the speed of a bee (relative to an obstacle) as well as the bee’s proximity to the obstacle. Thus, bees can use optic flow to gauge their distance from an obstacle [16,17], to reduce their flight speed as they approach an obstacle [20], to maintain their flight speed relative to an obstacle despite external wind [21], and to center themselves within a flight corridor, by moving laterally to balance optic flow across their left and right eyes [12,18,19]. In addition, the difference in optic flow produced by nearby obstacles versus the background helps bees gauge the dimensions and distance of obstacles [22,23]. Bees can enhance the visual information they receive from obstacles in the environment by performing side-to-side casting maneuvers as they fly or by slowing down and visually inspecting obstacles before continuing their flights, a behavior often associated with learning the layout of a new environment [22–24]. Overall, visual information is crucial for flying bees, and obstacles (e.g., cluttered vegetation) help provide the signals necessary for bees to successfully transit these structures. While visually guided bee flight is well-studied in simplified, artificial laboratory settings, we know little about how these principles operate in more complex, variable environments.

Cluttered vegetation also poses a flight hazard because collisions with vegetation can lead to irreparable wing damage [4], which impairs flight performance and is associated with higher mortality in bees [4,7]. To avoid collisions, bees flying through cluttered vegetation (or other obstacles) can use the visual information they gather to execute rapid lateral or vertical maneuvers, perform braking maneuvers, or take more sinuous paths around the obstacles [6,7,25]. Much of our empirical knowledge about obstacle traversal by bees is based on experiments in which bees are forced to transit simplified obstacles or apertures. Although these experiments reveal the physical mechanisms by which bees can traverse obstacles, they provide no information about how bees in nature negotiate complex physical environments, particularly given that bees have a variety of route choices available to them in natural settings, the most general of which are flying within (between) obstacles or bypassing obstacles entirely (e.g., flying above them). The choice to fly between obstacles may carry an increased risk of wing or body collisions [4], but it also provides strong visual signals that help bees control their ground speed and flight path (e.g., by centering themselves between lateral obstacles) [26]. Examining bee flight performance in the context of flight trajectory and route choice can help reveal how bees weigh the risk of collisions against the enhanced visual information and other potential benefits associated with flying through cluttered vegetation.

Bees flying in natural environments also regularly encounter wind, which can vary in speed, direction, and structure (e.g., periodic vortices, fully mixed turbulence), and each of these attributes of wind can affect flight performance in different ways. Bees flying into a steady wind can modify their flight speed relative to the air, to compensate for wind and maintain a constant speed relative to the ground [21]. However, headwinds containing periodic vortices (such as those shed behind a branch in wind) or fully mixed turbulence destabilize flying bees, impairing their ability to maintain a constant body orientation around the roll axis [5,27,28]. To compensate for this reduced roll stability, bees can increase the flapping frequency of their wings, modulate stroke amplitude to produce corrective asymmetries, and/or extend their legs, but many of these responses are likely to increase the energetic cost of flight [5,27]. Isolated gusts of wind can also cause flight instabilities, which trigger a suite of passive responses in bees such as altered body angle and flight speed, followed by active responses to return to their original body orientation and speed [29–31]. In addition, wind can increase the danger associated with some common flight maneuvers such as landing, with bees flying in wind tending to collide with landing surfaces rather than gradually slowing down to land [29,32]. Despite the growing body of research in this area, our knowledge about how wind affects flying bees remains limited to a fairly narrow set of experimental conditions, such as flight in open air streams without any physical clutter [5,27,31] or flight through vortices generated downstream of a single object in wind [28,29]. Thus, we know little about how wind affects the behavioral choices and flight performance of bees flying in more natural, cluttered habitats.

Bees encountering a large patch of cluttered vegetation in nature can choose to fly through the clutter or to fly above it, and they often make this choice while also contending with wind blowing in different directions relative to their flight path. To understand how wind, clutter, and route choice affect the flight performance of bees traversing complex environments, we filmed wild-caught honeybees (*Apis mellifera* Linnaeus 1758) flying through a laboratory enclosure containing a field composed of vertical obstacles. We varied obstacle field height (ranging from 11 to 127 mm tall) and wind condition (still air, headwinds or tailwinds). Obstacles in each field were arranged in longitudinal rows, providing two unobstructed flight corridors between the obstacles. The corridors between obstacles were 57 mm wide, which is ∼3X wider than a bee’s average wingspan [33]; thus, bees choosing to fly within the obstacles were not required to perform lateral maneuvers, but they did travel through a narrower corridor than those flying above the obstacles. We chose *A. mellifera* as a model organism because it is an important pollinator [34], its flight behavior is well-studied [17,33], and it shows little variation in body size [35], which helps eliminate one known source of variation in flight performance [6,7,36]. We reconstructed bees’ flight paths and used these data to answer two primary questions: (1) Is altitude affected by obstacle field height or wind condition? (2) Does altitude or wind condition affect flight performance?

## MATERIALS AND METHODS

### Experimental set-up

Experiments were conducted in a laboratory flight tunnel (20.0 × 19.1 × 115.0 cm; width x height x length) with a field of obstacles (hereafter referred to as the ‘obstacle field’) in the middle of the tunnel. The obstacle field consisted of vertical columns (diameter = 7 mm) arranged in three parallel rows of five obstacles each, running along the length of the tunnel (Figure 1a). Obstacles were made of dark green, cylindrical blocks (LEGO, Billund, Denmark) that contrasted with the black and white speckled pattern of the tunnel’s walls, and flight data indicated that bees were able to detect and avoid these obstacles. All obstacles within an obstacle field were of the same height, and the obstacles extended only partway to the tunnel’s ceiling, allowing bees to fly either within or above the obstacle field. The total height of the tunnel was 191 mm and the obstacle field heights tested were 11, 40, 69, 98, or 127 mm (Figure 1b), so a minimum of 64 mm (∼1/3 of the total vertical height) between the top of the obstacle field and the ceiling remained free of obstructions. We consider the 11-mm obstacle field as a control for the presence of obstacles because this obstacle field was too short for bees to fly within. There were approximately 20 mm between the outer rows of the obstacle field and the walls in each arrangement. Fans (AC Infinity, City of Industry, CA, USA) on each end of the tunnel produced a mild wind of the same mean flow speed (0.54 m s^-1^, measured with a Velocicalc Air Velocity Meter Model 9535, TSI, Shoreview, Minnesota, USA) above the obstacle field and within the corridors of the obstacle field (i.e., the space between the rows of obstacles); within the rows of obstacles (i.e., immediately downstream of an individual obstacle), the wind speed dropped to 0.36 m s^-1^ (Supplementary Figure S1).

**Figure 1.**
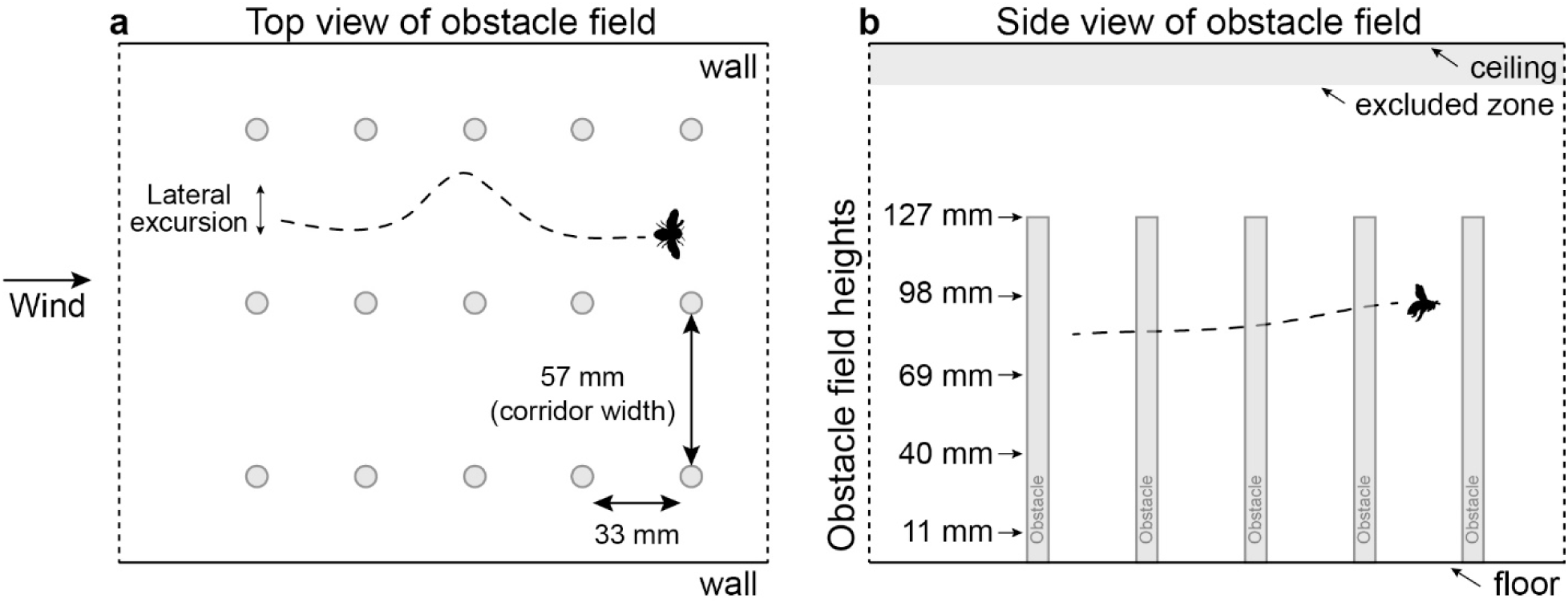
Schematic of experimental obstacle field. (a) Top view: obstacles in each field were arranged in three rows of five vertical columns each, forming two 57-mm wide, unobstructed flight corridors down the longitudinal axis of the enclosure. Bees’ flight paths within the horizontal plane (from above) were used to calculate lateral excursions (interquartile range of lateral positions throughout the flight). (b) Side view: the total height of the tunnel was 191 mm, and five obstacle field heights, ranging from 11 to 127 mm, were tested. Flights that crossed into the ‘excluded zone’ (within 15 mm of the ceiling; gray area on figure) were discarded.

Freely flying honeybees *Apis mellifera* (n = 58) were collected on the campus of the University of California, Davis. Single bees were flown in the tunnel with an obstacle field of one of the five possible heights, and the obstacle field height used for each bee was determined by a random number generator. Lights (26 Watts, full spectrum; Hagen, Mansfield, MA, USA) were alternately turned on and off at each end of the tunnel to motivate the bees to fly back and forth past the obstacle field (towards a light). We filmed between 5 and 13 flights per bee (mean = 9), with approximately half the flights in still air and half the flights with wind, for a total of 548 recorded flights. We randomly assigned each bee to begin their flights with either wind or still air. Bees flew in headwinds (flying into the wind) or tailwinds (flying with the wind), depending on the direction they flew relative to the air flow on a given transit. We define the two flight directions in our tunnel as ‘up-tunnel’ and ‘down-tunnel’ – bees experienced headwinds when flying in the up-tunnel direction with wind and tailwinds when flying in the down-tunnel direction with wind.

Flights were filmed with two synchronized Phantom v611 high-speed video cameras (Vision Research, Inc., Wayne, NJ, USA) sampling at 500 frames s^-1^, each positioned 30° from the vertical on opposing sides of the obstacle field and viewing down the length of the tunnel. Cameras were calibrated using a standard checkerboard calibration method and built-in MATLAB functions [37,38]. This method captures lens distortion and projective geometry (using the intrinsic parameters), as well as the global positions and orientations of the cameras relative to the flight tunnel (via the extrinsic parameters).

### Kinematic analysis

We used a detection and tracking pipeline to automatically track the centroid of bees in each camera view as they transited the obstacle field. From each frame, we subtracted the background and found one or more candidate positions of the bee using MATLAB’s built-in blob detection functions. We associated these detections into a single trajectory over time using a Kalman filter and Munkres’ assignment algorithm [39]. We then used DLTdv6 [40] to check and manually correct the automatically tracked positions of bees. We also labeled the positions of obstacles in the field using DLTdv6. Using the camera calibration, we converted the two-dimensional locations of the objects in each view into three-dimensional coordinates of the bees and obstacles. We analyzed bees’ trajectories from when they entered to when they exited the obstacle field (Figure 1a), and we smoothed the trajectories with quintic spline curves [41]. From each flight’s position data, we calculated the median altitude of the bee across its entire flight and the range of altitudes of the bee across its entire flight. To assess flight performance, we calculated two metrics: ground speed – the median of the bee’s speed relative to the ground, based on its movement in the horizontal plane (i.e., lateral and fore-aft motion), and lateral excursion, quantified by variation (i.e., interquartile range) in the bee’s lateral position over the entire flight.

### Statistical analyses

Because we captured wild bees outdoors, brought them into a novel setting (a laboratory flight enclosure with a fixed obstacle field height), and then recorded multiple flights by each bee, we first tested whether bees displayed any consistent changes in flight behavior (e.g., due to learning or familiarity) over the course of the flight trials. We compared flight data (altitude, ground speed, and lateral excursion) from each bee’s first recorded flight to that of its last recorded flight using paired Student’s *t*-tests, with separate analyses performed for flights in wind and still air, because the presence of wind covaried with flight number.

#### a. Is altitude affected by obstacle field height or wind condition?

We first tested whether median altitude and altitudinal range (maximum minus minimum altitude) of bees changed with experimental conditions. For these tests, we used the R function ‘lme’ from the package ‘nlme’ [42] to create linear mixed-effects models (LMM) with terms for wind (presence vs. absence), flight direction (up-vs. down-tunnel), and obstacle field height (5 categorical levels), and we allowed for interactions between these terms. Bee identities were included as a random effect to account for multiple observations per individual. Median altitude data were squared (*x*^2^) for normality and range data were log_10_-transformed for normality.

For the median altitude comparisons, data showed significantly different variances between experimental conditions (Levene’s tests, *P* < 0.005), which violated the homogeneity of variance assumption of the LMMs. To model unequal variances across experimental conditions, we used the R function ‘varIdent’ to update the original LMM with variance structures that were weighted for each experimental group – this resulted in seven new models corresponding to all possible combinations of equal/unequal variances across levels of wind, flight direction, and obstacle field height. These models, including the original LMM, were compared by their Akaike Information Criterion (AIC) using the R function ‘AIC’. The model with the lowest AIC, which compensated for unequal variances between obstacle field heights, was used for the main analysis in place of the original LMM. The standardized residuals of the final model showed similar variances between experimental groups (Levene’s test, *P* > 0.05 for significance) and appeared to be normally distributed (checked visually using quantile-quantile plots) [43].

#### b. Does altitude or wind condition affect flight performance?

We analyzed the two flight performance metrics (ground speed, lateral excursion) relative to altitude and wind condition. We defined altitude (using the median for each flight) relative to the obstacle field, in three ways: altitude above the tunnel floor and in the context of each obstacle field height (Altitude_floor_ * Obstacle height), altitude relative to the obstacle field (Altitude_obstacle_), and binomially, whether altitude was above or within the obstacle field (Route). These three definitions are closely correlated because (1) Altitude_obstacle_ = Altitude_floor_ - Obstacle height and (2) Route = “above the obstacles” if Altitude_obstacle_ > 0 and “within the obstacles” if Altitude_obstacle_ ≤ 0. To consider each of these definitions, we analyzed flight performance with three candidate LMM models that included an Altitude term based on each of these definitions. All models included terms for Wind and Direction, and bee identity was included as a random effect. Interactions were allowed between the Altitude, Wind, and Direction terms. Ground speed data were log_10_-transformed for normality, and lateral excursion data were cubic-root-transformed for normality. Assumptions of homogeneity of variances were checked with Levene’s test (*P* > 0.05 for significance) and assumptions of normality were checked visually using quantile-quantile plots [43]. For each flight performance metric, the candidate models were compared by AIC and the model with the lowest AIC was used to further analyze model terms. When applicable, multiple comparisons of model terms were done with Tukey Honest Significant Difference tests using the R package ‘lsmeans’ [44]. All statistical analyses were completed with R Statistical Software [45], using a critical *P-*value of 0.05 to determine statistical significance.

## RESULTS

Our analysis showed that the flight behavior of bees did not change in a consistent way over the course of the flight trials, in either still air or in wind (Student’s *t*-tests, *P* > 0.05; see Supplementary Table S1). Thus, we were able to treat the flight trials recorded from each individual as independent from one another.

### a. Is altitude affected by obstacle field height or wind condition?

Our analysis of altitudes revealed that trial conditions had only a minor effect on altitude.

Median altitude depended on obstacle field height (F_(4,53)_ = 5.795, *P* < 0.005), increasing in medians by 22.2 mm (23%) from the 11-mm obstacle fields to the 98-mm obstacle fields (df = 53, t-ratio = -3.634, *P* = 0.006) and 26.6 mm (28%) from the 11-mm obstacle fields to the 127-mm obstacle fields (df = 53, t-ratio = -3.881, *P* = 0.003) (Figure 2). Flight direction had a moderate effect on median altitude (F_(1,475)_ = 4.447, *P* = 0.036), but there were no significant pairwise differences in median altitude between the two flight directions (df = 475, t-ratio = - 1.018, *P* = 0.309) and no effect of wind on median altitude (F_(1,475)_ = 0.760, *P* = 0.384).

**Figure 2.**
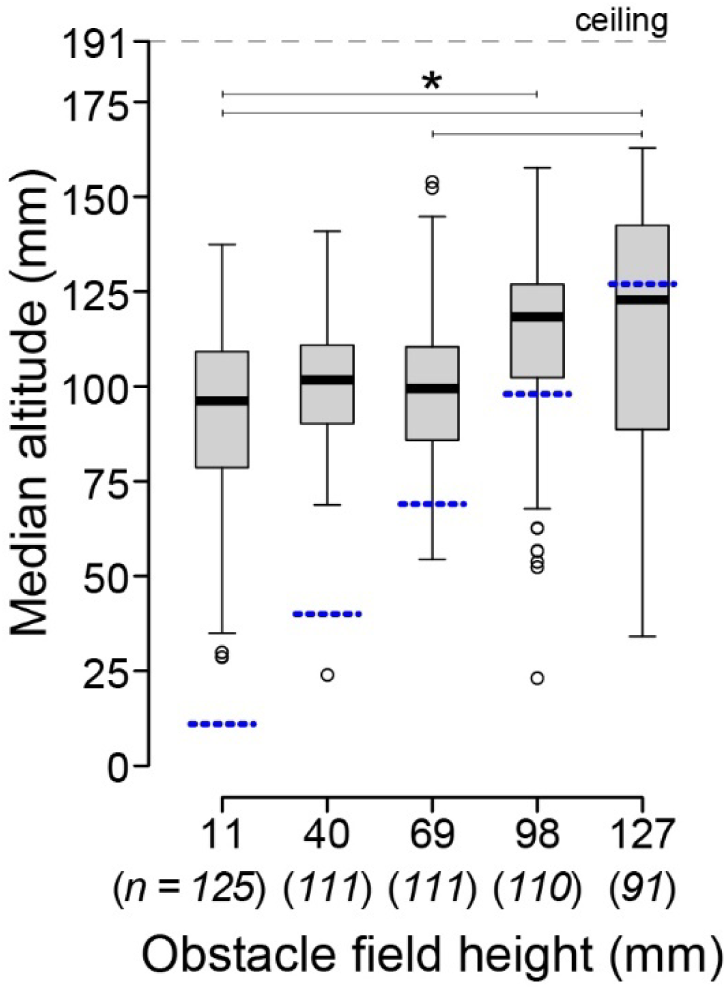
Median altitude of bees transiting obstacle fields. Dashed blue lines in indicate the obstacle field height relative to the flight altitude. Lines with asterisks indicate statistically different groups based on a linear mixed-effect model with Tukey post hoc comparisons (* *P* < 0.05). Italicized numbers below the x-axis labels show the sample size (number of flights) for each obstacle field height.

Bees maintained a relatively narrow range of altitudes as they transited the obstacle field (median range = 20.9 mm), and altitudinal range did not change with wind (F_(1,475)_ = 2.213, *P* = 0.138), flight direction (F_(1,475)_ = 0.356, *P* = 0.551), or obstacle field height (F_(4,53)_ = 2.535, *P* = 0.051). Overall, bees displayed similar altitudinal ranges across all conditions, and although there were a few statistically significant differences in median altitude, absolute differences in altitude were minor; for example, median altitudes increased in medians by only 26.6 mm between the shortest and tallest canopies, despite a 116-mm increase in obstacle field height (Figure 2).

### b. Does altitude or wind condition affect flight performance?

We tested whether ground speed and lateral excursions were affected by altitude, wind, and flight direction, using three definitions of altitude: median altitude relative to the top of the obstacle field, median altitude relative to the tunnel floor (in addition to a term for obstacle field height), and categorically, whether median altitude was above or within the obstacle field. Variation in ground speed was best explained by a model that used the categorical definition of altitude (above versus within the obstacle field; Figure 3). Ground speed was 40% faster above versus within the obstacle field (F_(1,483)_ = 19.542, *P* < 0.005), and it was 19% faster in tailwinds versus headwinds (df = 483, *t*-ratio = 3.590, *P* < 0.005) and 23% faster in tailwinds than in still air during down-tunnel flights (df = 483, *t*-ratio = 4.722, *P* < 0.005) (Figure 4).

**Figure 3.**
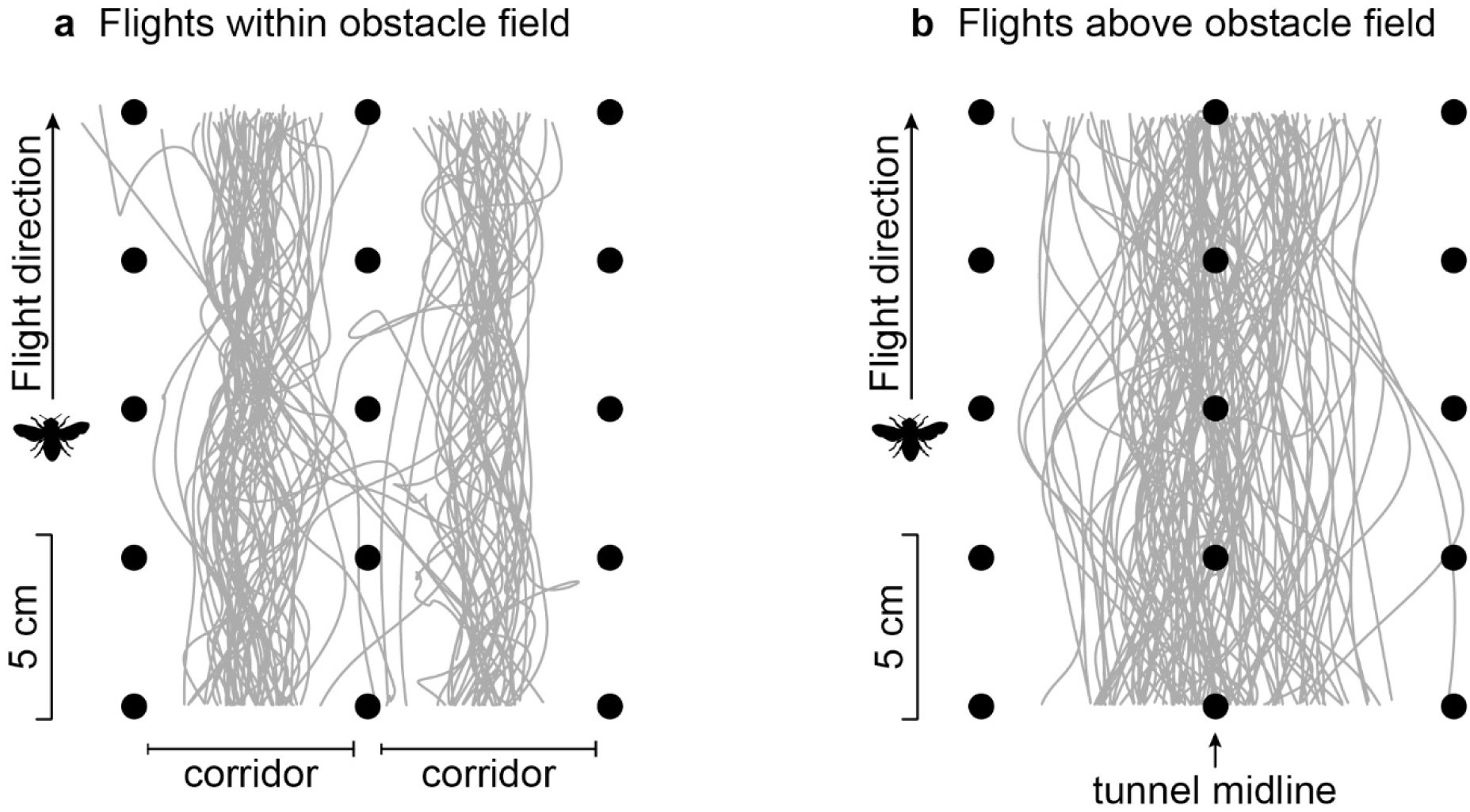
Top view of digitized flight trajectories for bees flying within (a) and above (b) the obstacle field. (a) Within the obstacle field, bees typically stayed within one of the two corridors formed by the rows of obstacles. All 82 flights within the obstacle field are shown. (b) Above the obstacle field, flights tended to be centered in the middle of the flight tunnel, above the middle row of obstacles. For clarity, traces of only 100 of the 466 flights above the obstacle field are shown.

**Figure 4.**
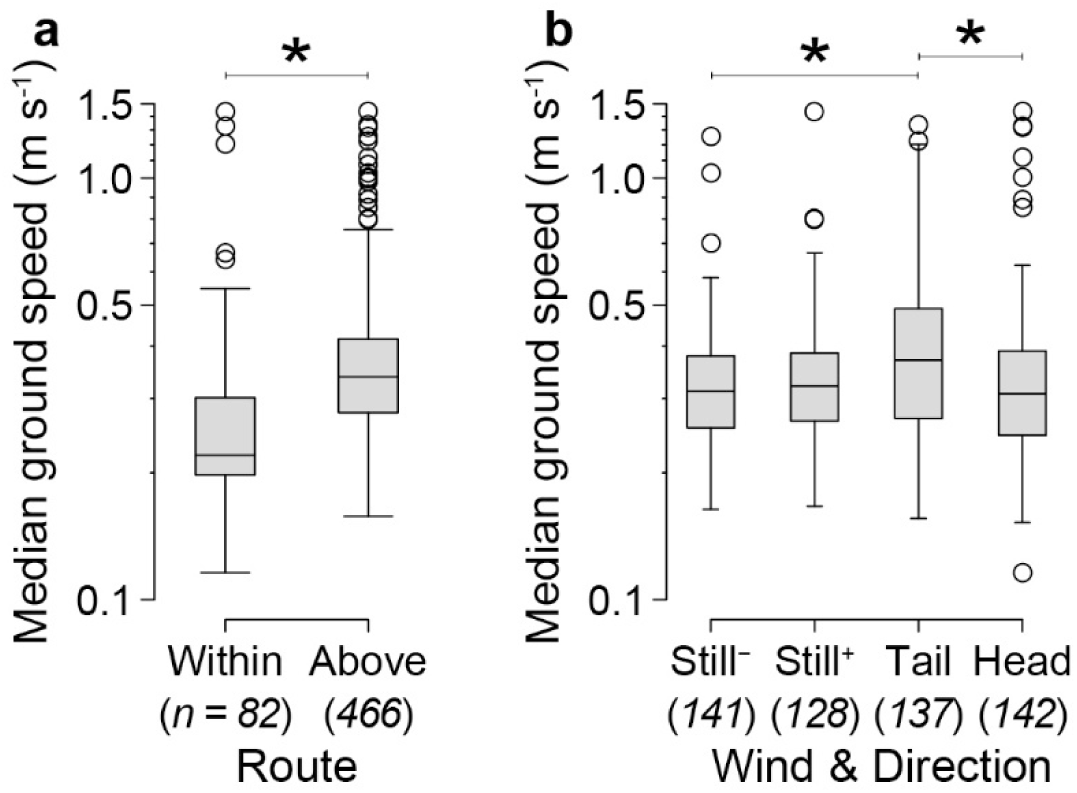
Effects of route and wind on ground speeds of bees transiting the obstacle field. (a) Median ground speeds were significantly higher above versus within the obstacle fields. (b) Median ground speeds were significantly higher in tailwinds than in headwinds and still air (down-tunnel direction only). “Still^-^” indicates down-tunnel flights in still air and “Still^+^” indicates up-tunnel flights in still air. Lines with asterisks indicate statistically different groups based on a linear mixed-effect model with Tukey post hoc comparisons (* *P* < 0.05). Italicized numbers below the x-axes labels show the sample size (number of flights) for each group.

Variation in lateral excursions was also best explained by a model that used the categorical definition of flight altitude. Lateral excursions were 36% larger above versus within the obstacle field (F_(1,483)_ = 7.096, *P* = 0.008) and 19% larger in wind than in still air (F_(1,483)_ = 12.888, *P* < 0.005) (Figure 5). Lateral excursions were not affected by flight direction (F_(1,483)_ = 1.685, *P* = 0.195).

**Figure 5.**
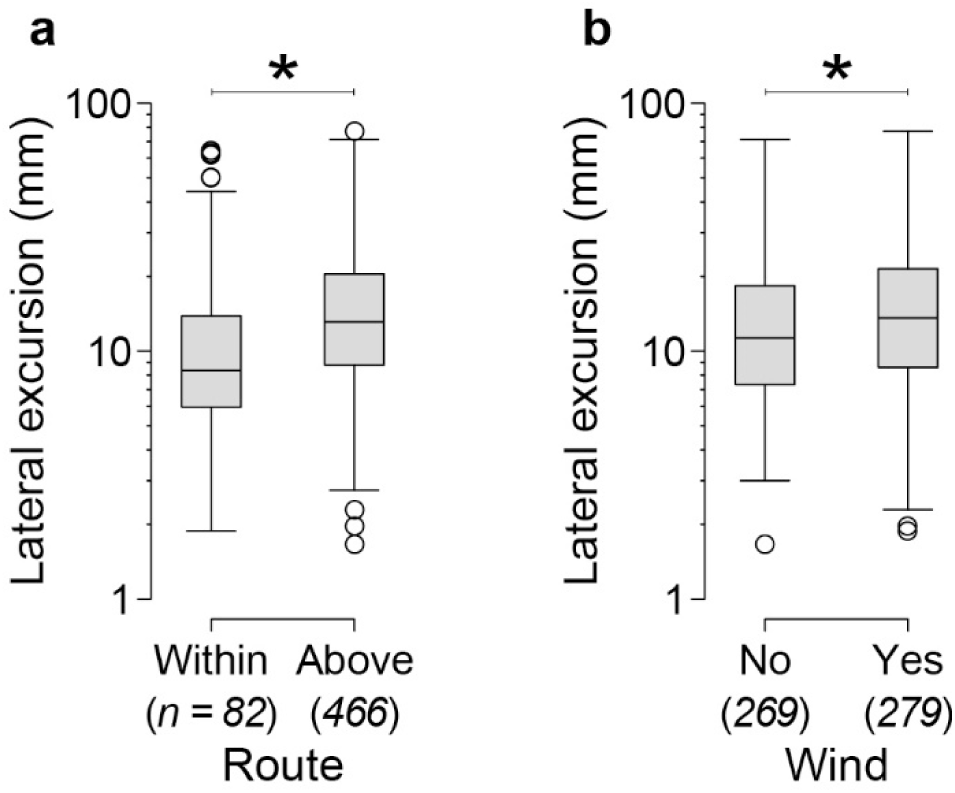
Effects of route and wind on lateral excursions of bees transiting the obstacle fields. Lateral excursions were (a) larger above versus within the obstacle fields and (b) larger in wind versus still air. Lines with asterisks indicate statistically different groups based on a linear mixed-effect model (* *P* < 0.05). Italicized numbers below the x-axes labels show the sample size (number of flights) for each group.

## DISCUSSION

### Is altitude affected by obstacle field height or wind condition?

Honeybee flights in the three shortest obstacle fields were concentrated around an altitude of approximately 100 mm, nearly halfway between the tunnel floor and the tunnel ceiling (Figure 2). However, when obstacle fields increased in height, bees increased their altitudes by approximately 20%. These results suggest that bees had a preferred altitude set by the dimensions of the flight tunnel (i.e., due to balancing optic flow from the walls, floor, and ceiling) and only responded to the obstacle field once it reached their preferred altitude. Furthermore, these results suggest that bees prefer to fly above obstacle fields, which is not surprising given that bees tend to fly routes that maximize the distance between their bodies and any surrounding landscape features [11,16,46]. This flight strategy can minimize the risk of collisions with obstacles, although in the present study only 7% of the 71 flights within the obstacle fields resulted in a collision. Thus, bees tended to adjust their altitudes higher to fly in the open space above the obstacle fields even though the obstacle field did not present a strong collision risk.

Altitude was not greatly affected by wind or flight direction, which was not surprising because the obstacle field did not significantly attenuate the wind speed [47] and the wind speed used in the experiment was not challenging compared to the wind in which honeybees are capable of flying [48]. In addition, if the obstacle fields were arranged in a manner that did reduce wind speeds (e.g., with staggered obstacles and/or closer obstacle spacing), the benefit of lower wind speeds within the obstacle field might be offset by the increased flight cost of maneuvering around obstacles and the increased risk of collision from flying closer to obstacles. More studies investigating bees’ use of vegetation and clutter as a refuge from wind are needed to fully understand how the animals weigh the risk of collisions and cost of maneuvering around obstacles against the altered wind conditions that the obstacles may offer.

### Do altitude or wind affect flight performance?

Variation in ground speed and lateral excursion were best explained by models that considered whether median altitudes were above versus within an obstacle field (Figure 3), but not necessarily how far the bees were above or below the obstacle fields or the tunnel floor. These results suggest that bees have a sharp transition in flight performance based on whether they are above versus within an obstacle field rather than a gradual transition in flight performance as they enter or exist an obstacle field.

Flights within the obstacle fields were characterized by reduced ground speeds and narrow lateral excursions (Figures 4, 5). Bees likely flew slowly within the obstacle fields as a way to balance the close proximity of the obstacles and maintain a preferred optic flow rate [12,16,49]. Although the bees could fly between obstacle corridors, most bees did not (Figure 3), suggesting that the narrow lateral excursions within the obstacle field were due to the reduced flight space in corridors. These data also suggest that the reduced ground speeds were not due to the bees attempting slowed, controlled turns around obstacles [6], which would have increased the magnitude of lateral excursions. Thus, the rapid transition in flight performance as bees lower into the obstacle field is likely due to the sudden visual feedback and narrow flight space provided by the obstacles in the field.

Wind caused significant shifts in flight performance for both ground speed and lateral excursion. Bees flew between 12 and 19% faster, based on medians, in tailwinds than in the three other wind and flight directions (Figure 4), suggesting that it was difficult for bees to adjust their air speed to maintain a preferred ground speed in tailwinds, when air is flowing from back to front and pushing the bee forward. For example, with a tailwind of 0.54 m s^-1^ and a preferred ground speed of 0.32 m s^-1^ (the median ground speed in still air), bees would need to fly *backwards* relative to the wind at approximately 0.22 m s^-1^ to maintain their preferred ground speed. However, the similarity in ground speeds between headwind and still air agrees with reports showing that insects, including honeybees, can maintain their preferred ground speed when flying in a headwind [21,50–53]. Thus, tailwinds present a unique flight challenge for honeybees within the context of our experimental set-up.

Wind increased the magnitude of lateral excursions, but this effect was likely limited to navigation rather than force production (e.g., air speed control) because, contrary to ground speeds, the effect occurred in both tailwinds and headwinds. Mechanical stimuli from wind can interfere with insect responses to visual stimuli [52], so the larger lateral excursions executed by honeybees in wind were possibly an intentional action to take in more visual information from the landscape to compensate for the mechanical challenge of wind [22,23]. To our knowledge, most studies on the effects of wind on bee flight have been conducted in headwinds [21,48], so additional studies that examine how steady winds from other directions interact with visual signals to affect flight performance are needed [54].

## Conclusions

Here we present data suggesting the vertical trajectories of bees are set by large-scale properties of the surrounding landscape, and that bees will only engage with clutter, such as vegetation, if it occupies their preferred flight space. Whether bees fly within the interstitial spaces of clutter, or avoid it entirely, can alter their flight performance: flights within clutter are slower and follow narrower paths, which are likely due to altered visual feedback and an increased risk of collisions. In contrast, wind can also alter flight performance (ground speeds are fastest in tailwinds; flight paths are wider in all windy conditions), but these effects may be due to inadequacies with flight control and processing visual versus mechanical stimuli from the environment. Thus, our small-scale laboratory experiment shows that flight challenges from the natural environment can impact bee behavior and flight mechanics for myriad reasons. Identifying the nuances of these effects and their underlying mechanisms can help us understand how honeybees in large-scale natural environments respond to similar types of flight challenges.

## ACKNOWLEDGEMENTS

This work was supported by the National Science Foundation (1711980 to N.P.B.).

## DATA ACCESSIBILITY

Data are available from the Dryad Digital Repository at https://doi.org/10.25338/B8Z34Q (Burnett et al., 2021).

