## Supplementary materials for "Wind and route choice affect performance of bees flying above versus within a cluttered obstacle field"

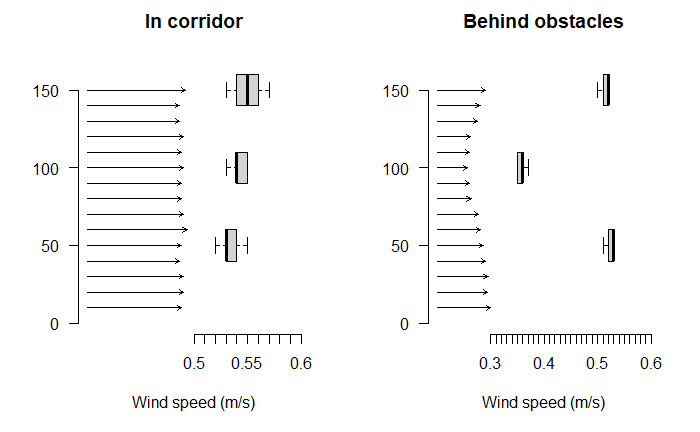
**Supplementary Materials**

**Figure S1.** Wind speeds in the experimental flight enclosure with a 127-mm tall obstacle field. Wind was measured in the middle of the obstacle field, halfway between the field’s entrance and exit, either in the corridor between the rows of obstacles (left) or directly downstream of the obstacles (right). Wind speed measurements were made with a Velocicalc Air Velocity Meter Model 9535 (TSI, Shoreview, Minessota, USA). The arrows in each panel show a vertical profile of wind speed, averaged over 20 seconds, with three replicate measurements per height. The boxplots show wind speeds (averaged over 20 seconds; 5 replicate measurements for each) for three heights. The highest measurement (150 mm) was above the 127-mm tall obstacle field and the other measurements were within the obstacle field. In the corridors, the wind speeds did not differ between these three heights (one way analysis of variance, F_(2,12)_ = 2.133, *P* = 0.161). Behind the obstacles, the wind speeds were faster above the obstacle field and near the base of the obstacles than they were just below the top of the obstacles (100 mm from the enclosure floor) (one way analysis of variance, F_(2,12)_ = 565.1, *P* < 0.005; Tukey Honest Significant Difference Test, *P* < 0.005).

| **Flight parameter** | **Still air** | **Wind** |
| --- | --- | --- |
| Median altitude | t = 0.903, df = 56, p = 0.370 | t = 0.921, df = 57, p = 0.361 |
| Altitudinal range | t = 0.065, df = 56, p = 0.948 | t = -0.052, df = 57, p = 0.959 |
| Median ground speed | t = -0.195, df = 56, p = 0.846 | t = -0.363, df = 57, p = 0.718 |
| Lateral excursion | t = -0.133, df = 56, p = 0.895 | t = -0.825, df = 57, p = 0.413 |

**Table S1.** Bees did not show a consistent change in their flight behavior over the course of the experiment. Results of paired *t­-*tests for the four flight parameters are shown, comparing the first recorded flight and the final recorded flight, with separate analyses for flights in still air and flights in wind.
